# Reevaluation of the Role of ERK3 in Perinatal Survival and Post-Natal Growth Using New Genetically-Engineered Mouse Models

**DOI:** 10.1101/467605

**Authors:** Mathilde Soulez, Marc K. Saba-El-Leil, Benjamin Turgeon, Simon Mathien, Philippe Coulombe, Sonia Klinger, Justine Rousseau, Kim Lévesque, Sylvain Meloche

## Abstract

The physiological functions of the atypical MAP kinase ERK3 remain poorly characterized. Previous analysis of mice with a targeted insertion of the *lacZ* reporter in the *Mapk6* locus (*Mapk6^lacZ^*) showed that inactivation of ERK3 in *Mapk6^lacZ^* mice leads to perinatal lethality associated with intrauterine growth restriction, defective lung maturation, and neuromuscular anomalies. To further explore the role of ERK3 in physiology and disease, we generated novel mouse models expressing a catalytically-inactive (*Mapk6^KD^*) or conditional (*Mapk6^Δ^*) allele of ERK3. Surprisingly, we found that mice devoid of ERK3 kinase activity or expression survive the perinatal period without any observable lung or neuromuscular phenotype. ERK3 mutant mice reached adulthood, were fertile and showed no apparent health problem. However, analysis of growth curves revealed that ERK3 kinase activity is ncessary for optimal post-natal growth. To gain insight into the genetic basis underlying the discrepancy in phenotypes of different *Mapk6* mutant mouse models, we analyzed the regulation of genes flanking the *Mapk6* locus by quantitative PCR. We found that expression of several *Mapk6* neighboring genes is deregulated in *Mapk6^lacZ^* mice, but not in *Mapk6^KD^* or *Mapk6^Δ^* mutant mice. Our genetic analysis suggests that off-target effects of the targeting construct on local gene expression are likely to be responsible for the perinatal lethality phenotype of *Mapk6^lacZ^* mutant mice.

## Introduction

Extracellular signal-regulated kinase 3 (ERK3) and ERK4 are atypical members of the mitogen-activated protein (MAP) kinase family (1). Despite their identification more than 25 years ago, very little is known about their physiological functions. The ERK3 gene (*MAPK6* in human) is expressed ubiquitously in adult mammalian tissues, while ERK4 gene (*MAPK4*) shows a restricted expression profile being detected mainly in brain tissue (2-4) (www.gtexportal.org/home). During mouse embryogenesis, ERK3 mRNA and protein levels peak around embryonic day 11, after the onset of organogenesis, and then decline thereafter (5). Expression of ERK3 is also up-regulated during *in vitro* differentiation of cell line models along the neuronal or muscle lineage (2, 6). These early observations have suggested a potential role for ERK3 signaling in cell differentiation and tissue development.

Analogous to classical MAP kinases, ERK3 is activated by phosphorylation of the activation loop, which stimulates its intrinsic catalytic activity and affinity for the substrate MK5 (7). Activation loop phosphorylation is mediated by Group I p21-activated kinases (8, 9), and is reversed by the action of the MAP kinase phosphatase DUSP2 (10). Of note, it has been previously proposed that ERK3 enzymatic activity is dispensable for MK5 activation and that ERK3 may exert a scaffolding function (11). However, other experiments using coupled *in vitro* kinase assays did not suppport this hypothesis (7). A more recent study reported that overexpression of ERK3 induces rounding of MDA-MB-231 breast cancer cells by an unknown mechanism independent of its kinase activity (12). The physiological importance of catalytic and non-catalytic functions of ERK3 remains an open question.

To investigate the physiological roles of ERK3, we have disrupted the *Mapk6* gene in the mouse by inserting a reporter *lacZ* gene in-frame with the ATG initiation codon in exon 2 (13). We found that loss of ERK3 function in *Mapk6^lacZ^* homozygous mice leads to intrauterine growth restriction, delayed lung maturation associated with defective type II pneumocyte differentiation, and neuromuscular anomalies. About 40% of *Mapk6^lacZ^* mutant mice died rapidly following delivery from respiratory distress syndrome. The other 60% of *Mapk6^lacZ^* pups survived the immediate neonatal interval but subsequently died within 24 hours from an unclear cause. These newborn mice were unable to feed and displayed symptoms of muscular weakness. All mice exhibited kyphosis and carpoptosis phenotypes. The perinatal lethality of *Mapk6^lacZ^* mutant mice has precluded the analysis of ERK3 functions in post-natal development and growth.

Here, we report on the generation of two novel genetically-engineered mouse models to address the importance of ERK3 catalytic activity and to study the role of ERK3 in post-natal development. We found that mice expressing a catalytically-inactive allele of ERK3 are born at normal Mendelian ratios and do not exhibit signs of respiratory distress or neuromuscular problems. Surprisingly, mice with a conditional disruption of the ERK3 gene also survived to adulthood, demonstrating that ERK3 expression and activity are dispensable for post-natal survival. However, we found that ERK3 catalytic activity is required for optimal post-natal growth in mice.

## Results and Discussion

### Mice expressing a catalytically-inactive ERK3 mutant survive to adulthood and are fertile

To address the physiological importance of ERK3 catalytic activity, we used a genetic approach to generate a mouse mutant bearing a kinase-dead (KD) allele of ERK3. To minimize disruption of the gene sequence, we have constructed a knock-in targeting vector that upon homologous recombination introduces the double mutation K49A/K50A in exon 2 of the *Mapk6* gene (Fig. 1A). Lys49 is part of the AXK motif in the β3 strand of the kinase domain N-lobe that bridges to the conserved Glu65 in the C-helix to anchor the α and β phosphates of ATP. The Lys-Glu salt bridge is considered a signature of the active kinase conformation and is essential for catalysis (14). To make sure that Lys50 could not substitute for Lys49, we also mutated this residue (Fig. 1B). As predicted, the K49A/K50A mutation abolishes ERK3 kinase activity *in vitro* (7). The targeting vector was electroporated in embryonic stem (ES) cells and two correctly targeted clones were identified by Southern blot analysis (Fig. 1C). One clone was injected in blastocysts to generate chimeric mice and was shown to transmit the *Mapk6^KD^* allele to the germ line.

**FIG 1.**
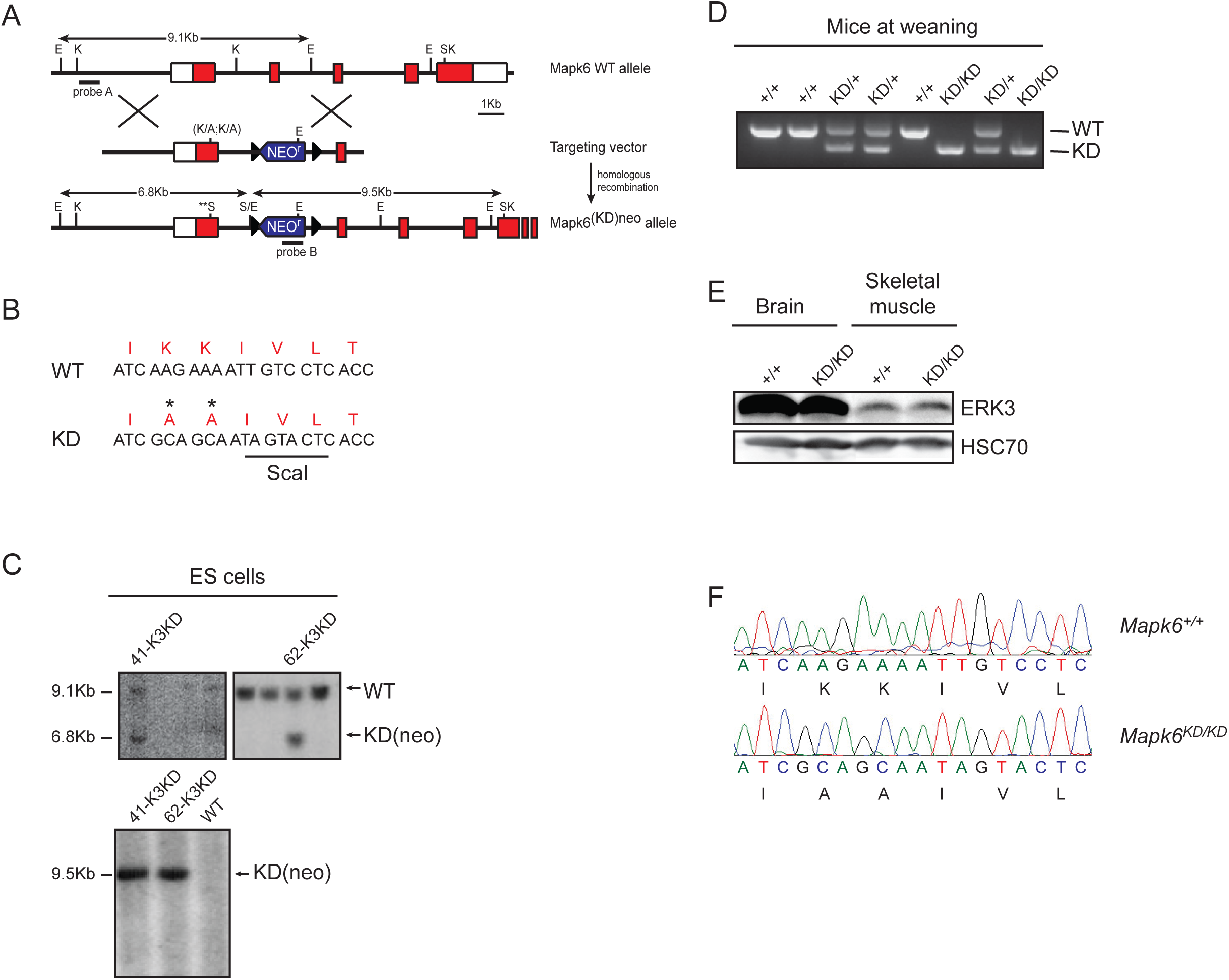
Generation of mice expressing kinase-dead ERK3. (A) Schematic representation of the gene targeting strategy used to generate a catalytically inactive knock-in allele of *Mapk6.* The targeting vector harbors a neomycin resistance gene (*neo^r^*) and DNA mutations that replace lysines 49 and 50 with alanine residues in exon 2 of the *Mapk6* coding sequence. Exons 2 to 6 and restriction sites are shown (E, EcoRI; K, KpnI; S, ScaI). Open boxes correspond to UTRs and closed boxes indicate coding regions. Black triangles correspond to loxP sites. (B) Nucleotide and amino acid sequences of codons 49 and 50 in the kinase domain of wild type and kinase-dead ERK3 protein. (C) Upper panel, Southern blot analysis of EcoRI-digested genomic DNA with probe A showing two correctly targeted ES clones with wild type (9.1 kb) and mutant (6.8 kb) alleles. Lower panel, Southern blot analysis of ScaI-digested genomic DNA with probe B showing the expected 9.5 kb recombinant band. (D) PCR analysis of a litter from a *Mapk6^KD/+^* heterozygous cross at weaning. (E) Immunoblot analysis of ERK3 protein expression in lysates prepared from brain and skeletal muscle of wild type and ERK3^KD/KD^ adult mice. (F) Chromatograms of genomic DNA sequences from *Mapk6^+/+^* and *Mapk6^KD/KD^* mice confirming the mutation of lysines 49 and 50 to alanines in exon 2 of the *Mapk6* gene.

Analysis of heterozygous *Mapk6^KD/+^* mice intercrosses revealed the presence of viable *Mapk6^KD/KD^* homozygous mice at the time of weaning (Fig. 1D). The mutant mice appeared healthy and showed no gross anatomical abnormality. Genotypes of offspring were distributed according to normal Mendelian inheritance (Table 1). *Mapk6^KD/KD^* mice survived to adulthood, were fertile, and remained generally healthy. We confirmed that ERK3 wild-type and KD proteins are expressed at similar levels in mouse tissues (Fig. 1E). Importantly, we further validated by DNA sequence analysis that *Mapk6^KD/KD^* mice express the *Mapk6^KD^* allele (Fig. 1F).

**Table1.** Genotypic analysis of offspring from *Mapk6^KD/^+* intercrosses^⋆^

| Stage | Total no. of pups | Genotype % |
| --- | --- | --- |
|  |  | +/+ : KD/+ : KD/KD |
| E18.5 | 136 | 27.2 : 47.8 : 25 |
| Adult | 192 | 25.5 : 53.7 : 20.8 |

\* Mixed C57Bl/6J-129/Sv genetic background

### ERK3 expression is dispensable for perinatal survival

The perinatal lethality of *Mapk6^lacZ^* mutant mice has precluded an analysis of the role of ERK3 in post-natal development and adult physiology. To circumvent this problem, we have generated a conditional allele of *Mapk6* by flanking exon 3 of the gene with *loxP* sites (Fig. 2A). Two correctly targeted ES clone were identified (Fig. 2B) from which a single clone produced chimeric mice that could transmit the floxed allele to the germ line. Deletion of exon 3 in the kinase domain is predicted to disrupt the kinase fold and introduce a frameshift to yield a null allele. To validate that we had created a conditional null allele of *Mapk6*, we crossed *Mapk6^flox/flox^* mice to *Sox2*-*Cre* transgenic mice to inactivate *Mapk6* in the epiblast and resulting embryo. Immunoblot analysis of E12.5 embryos confirmed the absence of ERK3 protein in *Mapk6^Δ/Δ^* (Δ symbolizes the Cre-excised allele) homozygous embryos (Fig. 2C).

**FIG 2.**
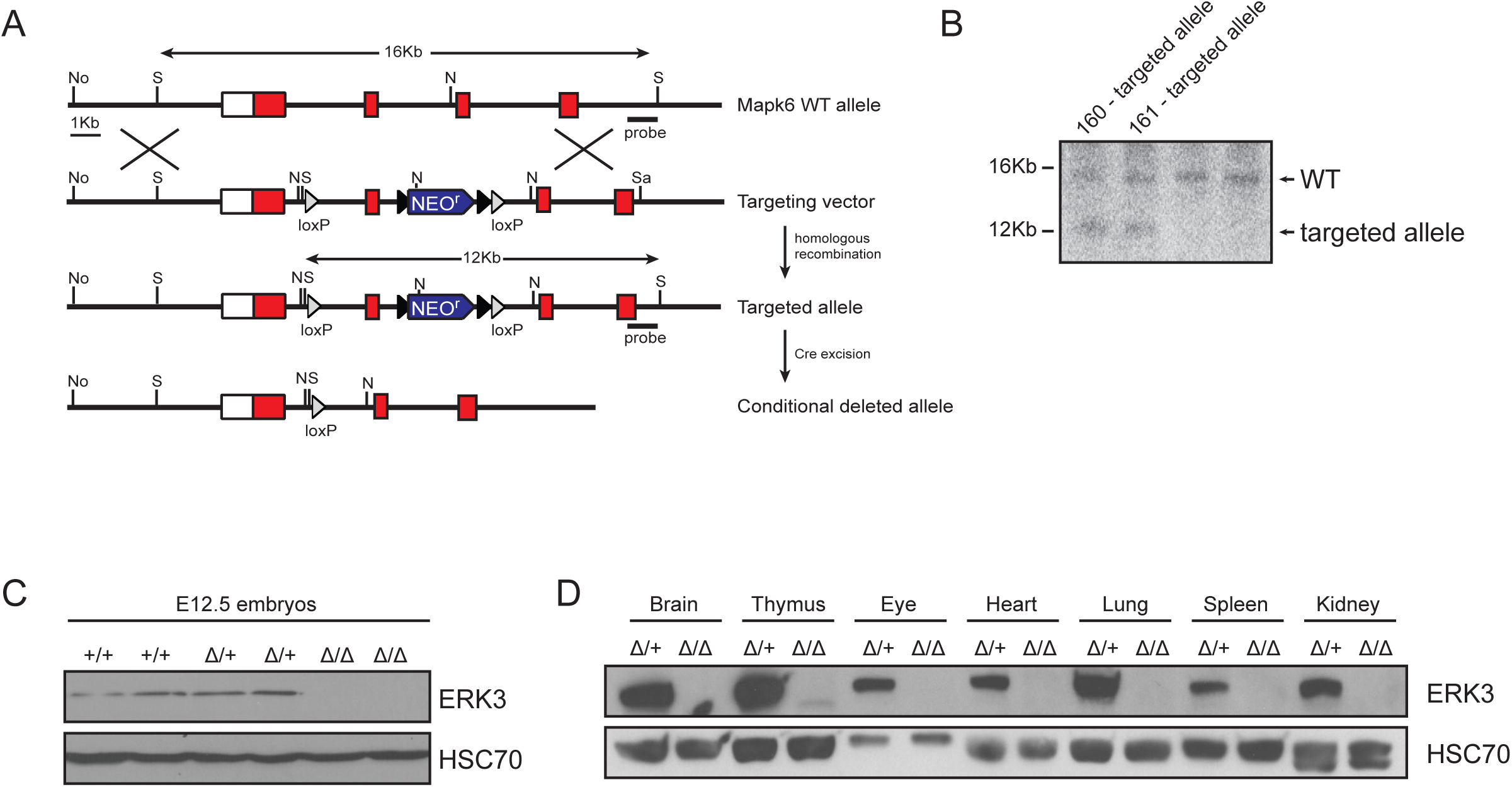
Generation of mice expressing a conditional *Mapk6* allele. (A) Schematic representation of the *Mapk6* locus, targeting vector, recombinant conditional allele and conditional deleted allele. The targeting vector carries a neomycin resistance cassette (*neo^r^*) flanked by two FRT sites and exon 3 is flanked by two loxP sites 3. Exons 2 to 5 and restriction sites are shown (N, NheI; N, NotI; S, Sfil). Open boxes correspond to UTRs and closed boxes indicate coding regions. Whitey and black triangles correspond to loxP and FRT sites, respectively. (B) Southern blot analysis of Sfil-digested genomic DNA showing two correctly targeted ES clones. Wild type (16 kb) and targeted mutant (12 kb) alleles are indicated. (C) Immunoblot analysis of ERK3 protein expression in lysates prepared from wild type (+/+), heterozygous *Mapk6^Δ/+^ (Δ/+*) and *Mapk6^Δ/Δ^* (*Δ/Δ*) E12.5 embryos. (D) Immunoblot analysis of ERK3 protein expression in lysates prepared from different tissues of *Mapk6^Δ/+^* and *Mapk6^Δ/Δ^* adult mice.

To verify that the Cre-excised *Mapk6^μ^* allele behaves like the *Mapk6^lacZ^* null allele, we intercrossed *Mapk6^Δ/+^* mice. Surprisingly, we observed that ERK3-deficient *Mapk6^Δ/Δ^* mice are born at the expected Mendelian frequency (Table 2) and survive to adulthood without any apparent health issues. Immunoblot analysis of various tissues isolated from adult*Mapk6^Δ/Δ^* mice confirmed the absence of ERK3 expression in all tissues examined (Fig. 2D). *Mapk6^Δ/Δ^* mice were further shown to be fertile. We conclude that ERK3 expression is dispensable to survive the perinatal period.

**Table2.** Genotypic analysis of offspring from *Mapk6^Δ/+^* intercrosses^⋆^

| Stage | Total no. of pups | Genotype % |
| --- | --- | --- |
|  |  | +/+ : Δ/+ : Δ/Δ |
| E18.5 | 90 | 35.6 : 50 : 14.4 |
| Adult | 110 | 23.6 : 55.5 : 20.9 |

\* Mixed C57Bl/6J-129/Sv genetic background

### Phenotypic analysis of *Mapk6^KD/KD^* and *Mapk6^Δ/Δ^* mutant mice

Homozygous *Mapk6^lacZ^* mutant mice display an intrauterine growth restriction phenotype, which is associated with adverse perinatal outcome (13). In contrast, we did not observe any decrease in the body weight of heterozygous or homozygous *Mapk6^KD^* and *Mapk6^Δ^* E18.5 embryos as compared to wild type littermates, consistent with their perinatal survival (Fig. 3A, B and C). We also reported that *Mapk6^lacZ/lacZ^* mutant mice exhibit a lung maturation defect characterized by decreased saccular space and the persistence of type II pneumocytes with high glycogen content (13). Histological examination of the lungs of *Mapk6^KD/KD^* and *Mapk6^Δ/Δ^* mutant E18.5 embryos revealed a normal lung architecture with the saccular space comparable to wild type animals (Fig. 3D). *Mapk6^KD/KD^* and *Mapk6^Δ/Δ^* mutant E18.5 embryos appeared morphologically normal and did not exhibit the smaller size and carpoptosis (wrist drop) phenotypes of *Mapk6^lacZ/lacZ^* mutant mice (Fig. 3E). These results argue that the phenotypes of *Mapk6^lacZ^* homozygous mutant mice originally reported are not a direct consequence of the loss of ERK3 activity or expression but may rather reflect an off-target effect of the targeting construct.

**FIG 3.**
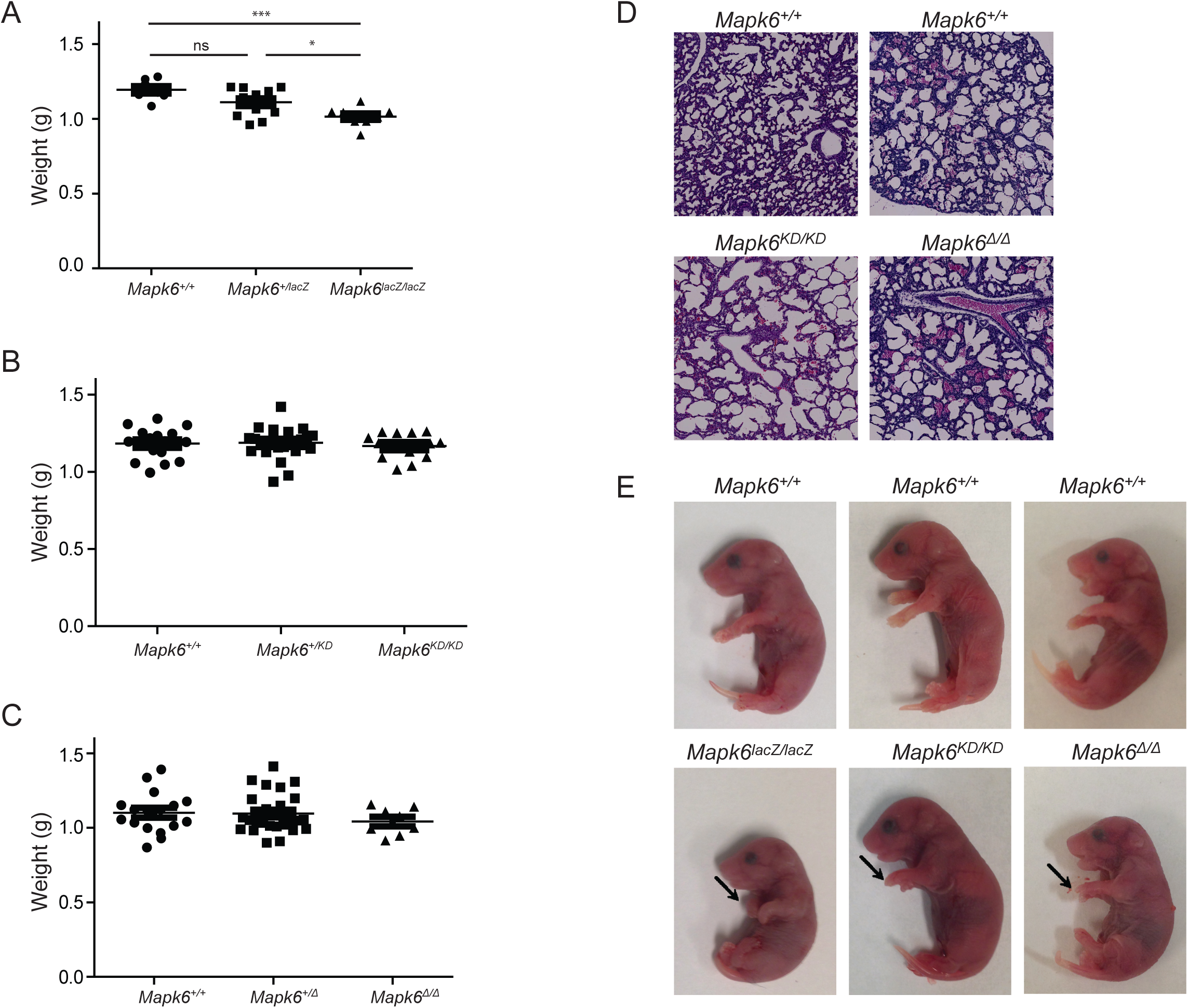
Phenotypic analysis of *Mapk6^KD/KD^* and *Mapk6^Δ/Δ^* E18.5 embryos. (A-C) Body weight distribution of E18.5 embryos from *Mapk6^lacZ/+^* (A), *Mapk6^KD/+^* (B) and *Mapk6^Δ/+^* intercrosses. ^∗^ p < 0.05, ^∗∗∗^ p < 0.001 (unpaired Student’s *t*-test). (D) Representative photographs of H/E staining of lung sections from E18.5 embryos of the indicated wild type or mutant *Mapk6* genotype. (E) Representative images of E18.5 embryos of the indicated wild type or mutant *Mapk6* genotype. Arrows point to the wrists and show carpoptosis which is present exclusively in *Mapk6^lacZ/lacZ^* embryos.

### Targeted insertion of the *Mapk6^lacZ^* construct influences neighboring gene expression

Insertion of foreign DNA fragments into the genome may have unintended effects on the expression and function of neighboring genes. In a recent study, West et al. (15) have used RNA-seq to analyze in a systematic manner the transcriptional impact of targeted mouse mutations on local changes in gene expression. They measured the expression levels of genes within +/− 500 kb of the targeted gene in tissues from 44 homozygous mutants generated by the International Knockout Mouse Consortium (16) compared with C57BL/6N inbred control mice. The targeting strategies insert a bacterial beta-galactosidase reporter (*lacZ* gene) downstream of the target gene endogenous promoter and a neomycin resistance selection cassette, similar to the strategy used to produce *Mapk6^lacZ^* mutant mice. The authors found that one or more neighboring genes is down-regulated or up-regulated in up to 59% of the mouse mutants, depending on the targeting approach (15). Various mechanisms have been proposed to explain this local transcriptional dysregulation, including disruption of regulatory and insulating elements with cis-acting consequences and effects of the exogenous *neo^r^* promoter on expression of 3’ genes.

The impact of local changes in gene expression on the phenotypic spectrum of targeted mouse mutants has not been thoroughly studied. However, an accumulating number of reports indicate that some phenotypes associated with a targeted gene mutation are causally related to off-target effects on neighboring genes (17-21). For example, mice with a homozygous disruption of the *Ampd1* gene generated by a targeted trap mutation (knockout-first allele) die within two days post-natally, and show reduced body weight, decreased lung saccular space and lack of milk in their stomach (20). However, after removal of the knockout-first cassette containing the *lacz* and *neo^r^* genes to generate a conditional allele and further deletion of exon 3 of the *Ampd1* gene to produce a null allele, homozygous mutant mice survived to adulthood. RNA-seq analysis revealed that expression of the neighboring *Man1a2* and *Nras* genes was down-regulated in the knockout-first *Ampd1* mice, but not in the conditional *Ampd1* mutant mice. Another recent study reported that replacement of the coding region of the steroid metabolising enzyme gene *Hsd17b1* with the *lacZ* gene inserted in the translation initiation site and a *neo^r^* selection cassette (Hsd17b1-LacZ/Neo mice) results in reduced adipose mass, increased lean mass and lipid accumulation in the liver of male mice (17). Further characterization revealed that the metabolic phenotype of Hsd17b1-LacZ/Neo mice is caused by the reduced expression of the immediate 5’ gene *Naglu*, which encodes the N-acetyl-alpha-glucosaminidase enzyme, and not by a deficiency in HSD17B1 activity. Local changes in gene expression due to vector insertion may also explain the differences in phenotypes observed for multiple null mutants of the same gene (19).

These considerations led us to evaluate the potential impact of the targeted *Mapk6^lacZ^* mutation on the transcription of neighboring genes. We first focused on genes within plus or minus 1 Mb of the *Mapk6* locus on mouse chromosome 9 (Fig. 4A) and quantitatively measured their level of expression by qRT-PCR in littermate wild type and *Mapk6^lacZ^* homozygous E12.5 embryos. Of the 18 genes found in this interval, 15 were found to be significantly expressed. We observed a global dysregulation of the expression of *Mapk6* neighboring genes in *Mapk6^lacZ/lacZ^* mutant embryos, with 7 genes being statistically up-regulated by more that 50% (Fig.4B). In contrast, none of the genes was deregulated in *Mapk6^KD/KD^* (Fig. 4C) or *Mapk6^Δ/Δ^* (Fig. 4D) embryos as compared to their respective wild type littermate controls. We then selected 14 genes associated with embryonic development outside the 1 Mb interval around *Mapk6* to determine whether the *Mapk6^lacZ^* mutation exerts longer-range transcriptional effects. Notably, we observed that the expression of 5 genes was significantly up-regulated in *Mapk6^lacZ/lacZ^* embryos as compared to littermate *Mapk6^+/+^* embryos (Fig. 4E), whereas none of the genes were deregulated in *Mapk6^KD/KD^* (Fig. 4F) or *Mapk6^Δ/Δ^* (Fig.4G) mutant embryos. These results indicate that targeted insertion of the *Mapk6^lacZ^* construct exerts unintended effects on the regulation of local gene expression.

**FIG 4.**
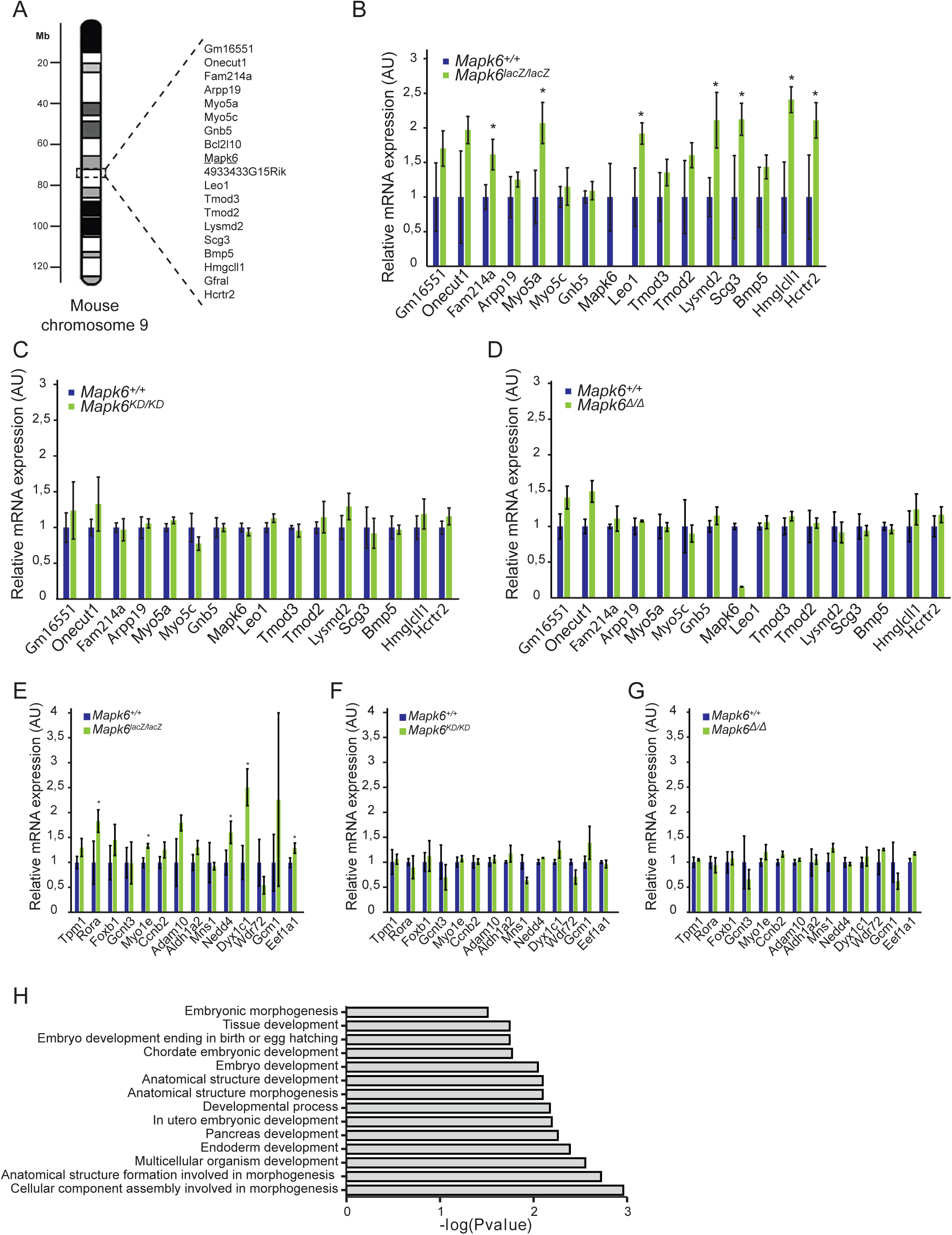
Targeted insertion of the *Mapk6^lacZ^* construct influences the transcription of neighboring genes. (A) Schematic representation of mouse chromosome 9 and neighboring genes of *Mapk6* within a 1Mb interval. (B-D) Relative mRNA expression of 15 genes within +/− 1 Mb interval around *Mapk6* in E12.5 embryos of wild type or homozygous *Mapk6^lacZ/lacZ^* (B), *Mapk6 ^KD/KD^* (C) and *Mapk6^//^* (D) genotypes. Gene expression was measured by quantitative RT-PCR. ^⋆^ p < 0.05 (unpaired Student’s *t*-test). (E-G) Relative mRNA expression of 14 developmental genes on mouse chromosome 9 outside 1 Mb from the *Mapk6* locus in E12.5 embryos of wild type or homozygous *Mapk6^lacZ/lacZ^* (E), *Mapk6 ^KD/KD^* (F) and *Mapk6^Δ/Δ^* (G) genotypes. ^∗^ p < 0.05 (unpaired Student’s *t*-test). (H) GO term enrichment analysis of the 89 genes located within +/− 5 Mb of the *Mapk6* locus on mouse chromosome 9 using the online software DAVID. Enrichment of Biological Processes is expressed as log10 p-value.

We next performed Gene Ontology (GO) enrichment analysis (22) of the 89 genes within plus or minus 5 Mb of the *Mapk6* locus using the online software DAVID (23). Interestingly, 14 out of 88 significantly enriched Biological Process GO term (defined as GO term for which Fisher exact p-value < 0.05) were associated with embryogenesis, morphogenesis or development, revealing an enrichment of genes involved in developmental processes in the 10 Mb chromosomal region surrounding the *Mapk6* locus (Fig. 4H). We hypothesize that the off-target
effects of the *Mapk6^lacZ^* construct are responsible for the perinatal lethality phenotype of *Mapk6^lacZ^* mutant mice rather than the loss of ERK3 function.

### ERK3 activity is required for optimal post-natal growth in mice

The availability of surviving *Mapk6* mutant mice allows to explore the function of ERK3 in post-natal development. Despite the fact that *Mapk6^KD/KD^* and *Mapk6^Δ/Δ^* mutant mice have a comparable body weight to wild type mice at birth, we noticed that some mutant mice appear smaller at weaning. To investigate the possible role of ERK3 in post-natal growth, we measured the weight of mouse cohorts of different *Mapk6* genotypes from birth through weaning at 3 weeks of age. The growth rate of *Mapk6^Δ/Δ^* homozygous mice was clearly reduced as compared to control *Mapk6^flox/+^* mice and heterozygous *Mapk6^Δ/flox^* or *Mapk6^Δ/+^* mice (Fig. 5A). The difference in body weight already reached statistical significance at day 5 and was maintained up to the time of weaning (Fig. 5A and B). A similar phenotype was observed for *Mapk6^KD/KD^* mice (Fig. 5C and D). These results provide strong evidence that ERK3 kinase activity is necessary for optimal post-natal growth in mice. Future studies will be required to understand the cellular and molecular basis of this complex phenotype, and to evaluate its long-term physiopathological impact.

**FIG 5.**
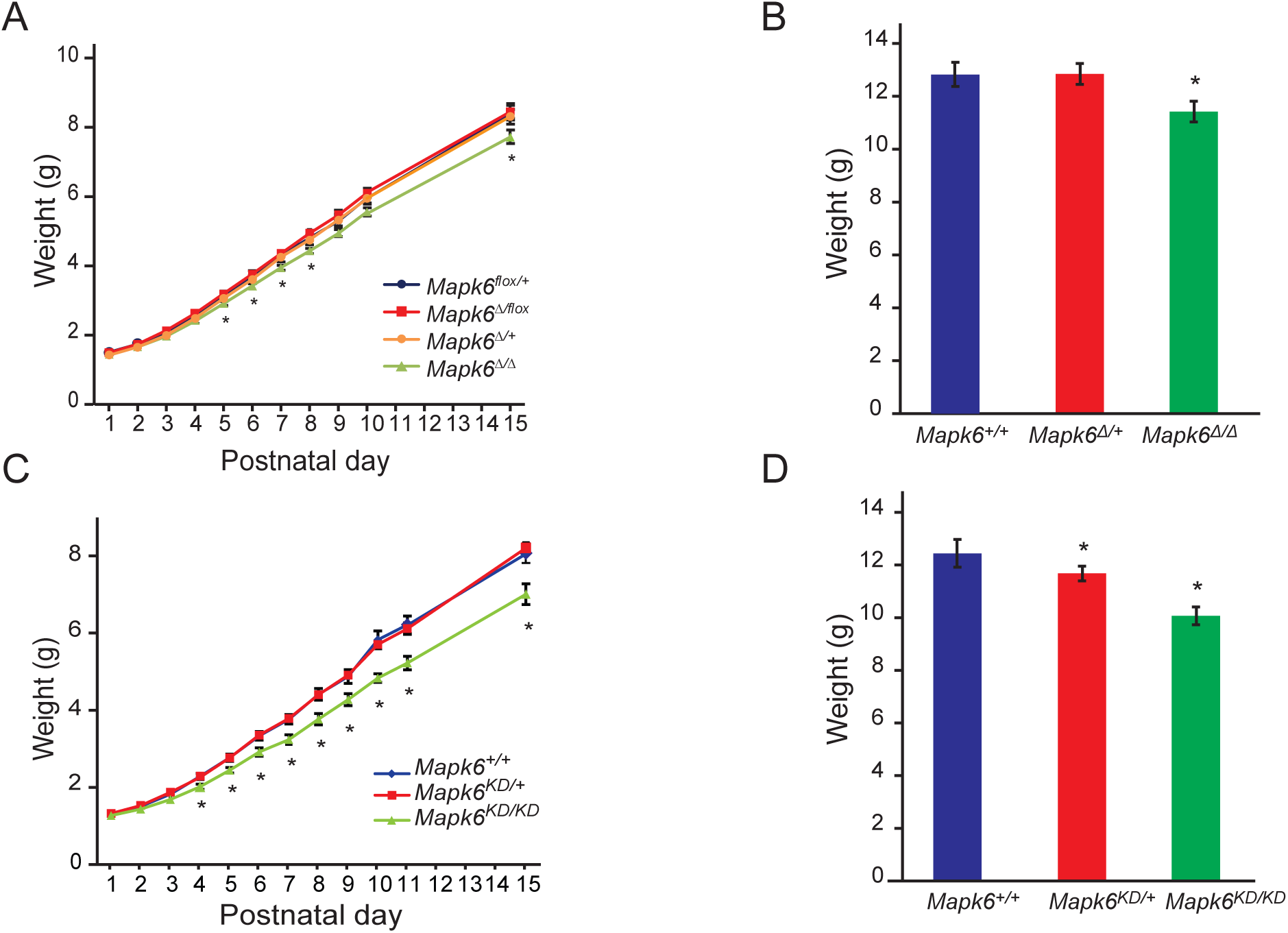
Impact of the loss of ERK3 activity or expression on post-natal growth in mice. (A) Growth curves of *Mapk6^flox/+^*, *Mapk6^Δ/flox^, Mapk6^Δ/+^* and *Mapk6^Δ/Δ^* mice (n ≥ 11) over 15 postnatal days. (B) Analysis of *Mapk6^+/+^*, *Mapk6^Δ/+^* and *Mapk6^Δ/+^* mice (n ≥ 13) body weight at weaning. (C) Growth curves of *Mapk6^+/+^*, *Mapk6^KD/+^* and *Mapk6^KD/KD^* mice (n ≥ 12) over 15 post-natal days. (D) Analysis of *Mapk6^+/+^, Mapk6^KD/+^* and *Mapk6^KD/KD^* mice (n ≥ 19) body weight at weaning (day 21). ^∗^ p < 0.05 (unpaired Student’s *t*-test).

### Conclusions

We have generated two novel genetically-engineered models of *Mapk6* mutant mice. We show that ERK3 activity and expression are dispensable for perinatal survival, contrary to previous findings obtained from the analysis of *Mapk6^lacZ^* mice. We also unveil a novel role of ERK3 kinase activity in regulating post-natal growth. These mouse models will provide valuable tools to study the poorly understood physiological functions of the atypical MAP kinase ERK3. The *Mapk6^KD^* mutant mouse will also be useful to genetically validate the selectivity of future small molecule inhibitors of ERK3. Our findings also emphasize the importance of evaluating the impact of targeted mutant constructs on local gene regulation and, whenever possible, to analyze more than one mouse line harboring different mutant alleles for interpreting the phenotypes associated with mutations in a specific gene.

## Materials and Methods

### Generation of mouse mutants

To generate mice bearing a catalytically inactive allele of *Mapk6*, we introduced a mutation in the essential AXK motif of the kinase domain in exon 2 by homologous recombination. Part of exon 2 was amplified by PCR using primers containing a modified DNA sequence to change the sequence coding for lysines 49 and 50 (AAG AAA) by a sequence coding for two alanine residues (GCA GCA). A ScaI restriction site was also introduced to reassemble the mutated exon 2 in a pBluescript II cloning vector. Left and right arms of genomic DNA coding for intronic sequences were PCR amplified and cloned upstream and downstream of the mutated exon 2. A second right arm containing exon 3 was added downstream of the shorter right arm allowing to insert an XhoI restriction site in which the *neo^R^* selection cassette flanked by *loxP* sites was introduced. The targeting vector was linearized and electroporated into G4 mouse ES cells. Recombinant clones were selected with G418 and screened by Southern blot analysis for homologous recombination. One correctly targeted ES cell clone was injected into C57BL/6 blastocyst-stage embryos to produce chimeric mice, and was shown to transmit the mutant allele to the germ line.

We generated a conditional *Mapk6* allele by flanking the essential exon 3, which encodes kinase subdomains VIII and IX, by *loxP* sites. Briefly, this was achieved by assembling left and right arm genomic fragments containing exon 2 and exons 4 and 5, respectively, with exon 3 by Gibson Assembly (New England Biolabs). An upstream *loxP* site was inserted in intron 2 upstream of exon 3 and an FRT-flanked *neo^R^* selection cassette followed by a second *loxP* site was inserted downstream of the exon. The targeting vector was linearized and electroporated into G4 mouse ES cells. Recombinant clones were screened as described above. A correctly targeted ES cell clone was injected into C57BL/6 blastocyst-stage embryos to produce chimeric mice, and germline transmission was obtained.

### Genotyping and DNA sequencing

Genomic DNA was extracted using phenol/chloroform protocol and was subjected to PCR. *Mapk6^KD/KD^* genotyping was performed with the following priming sequences: wild-type and kinase-dead allele CTG TTT CTG CCT CCC ATG TG; wild-type allele GAT AGA TTC ATC TAT CTC TCC C: kinase-dead allele ATT CCA TCA GAA GCT ATA AAC TTC G. For sequencing of the kinase dead allele a PCR reaction amplifying exon 2 of *Mapk6* (priming sequences GCT TTA AGT CTG AGG GGA AC and GCA GAA TGG ATA TAT TTG AG) was first perform followed by Sanger sequencing using DNA analyser ABI 3730. *Mapk6^Δ/Δ^* genotyping was performed with the following priming sequences: Δ allele CCT GCC TCC AAA CTG ATA CTG and GAC GTG ATC CAC CTA ATA ACT TCG; exon 3 (wild-type and flox) AAC ACT GAC CTG CAA AGA GG and TCT CTC CCT CAC CTT TCC C; flox allele ACC ACC AAG CGA AAC ATC G and GAC GTG ATC CAC CTA ATA ACT TCG; wild-type allele TGC AGG TCA GTG TTG TAT GG and AGT GCT TGA TTT GAA ACA CAG G.

### Animal husbandry and experimentation

Animals were housed under pathogen-free conditions and were handled according to procedures and protocols approved by the Université de Montréal Institutional Animal Care Committee. Male and female mice maintained on mixed C57BL/6/129/Sv genetic background were used for these studies. For timed matings, pregnant females were sacrificed at gestation day 12.5 or 18.5 by CO2 euthanasia and embryos were removed by caesarean section. Each embryo was weighted and processed for further analysis.

### Histology

Lungs were fixed in 10% formalin, embedded in paraffin, and sliced in 5-μm thin sections. Tissue sections were mounted on glass slides and stained with hematoxylin/eosin (H/E) using a standard protocol.

### Immunoblot analysis

Cell lysis and immunoblotting analysis were performed as described (6) using commercial anti-ERK3 (Abcam, dilution 1:1000) and anti-HSC70 (Santa Cruz Biotechnology, dilution 1:2000) antibodies.

### Quantitative RT-PCR

Total RNA was isolated from E12.5 embryos, purified with the RNeasy kit (Qiagen) and reverse transcribed using the Maxima First Strand cDNA synthesis kit with ds DNase (ThermoFisher Scientific). Gene expression was determined using assays designed with the Universal Probe Library from Roche (www.universalprobelibrary.com). For each qPCR assay, a standard curve was performed to ensure that the efficiency of the assay was between 90% and 110%. Target DNA amplification was measured on the Viia7 Real-Time PCR System (Life Technologies) programmed with an initial step of 20 sec at 95°C, followed by 40 cycles of: 1 sec at 95°C and 20 sec at 60°C. Relative expression (RQ = 2^−ΔΔCT^) was calculated using the ExpressionSuite software (Life Technologies), and normalization was done using both GAPDH and ACTB.

## Acknowledgments

We thank Mélania Gombos for help with animal experimentation, Sébastien Harton for technical support in generating the *Mapk6^flox^* mouse line, and Raphaelle Lambert for qPCR analysis. This work was supported in part by grants from the Canadian Institutes for Health Research to S. Meloche. S. Meloche held the Canada Research Chair in Cellular Signaling.

## References

1. Mathien S, Soulez M, Klinger S, Meloche S. 2018. Erk3 and Erk4, p 1632–1638. In Choi S (ed), Encyclopedia of Signaling Molecules, Second Edition ed. Springer International Publishing, Cham, Switzerland.

2. Boulton TG, Nye SH, Robbins DJ, Ip NY, Radziejewska E, Morgenbesser SD, DePinho RA, Panayotatos N, Cobb MH, Yancopoulos GD. 1991. ERKs: a family of protein-serine/threonine kinases that are activated and tyrosine phosphorylated in response to insulin and NGF. Cell 65:663–675.

3. Turgeon B, Saba-El-Leil MK, Meloche S. 2000. Cloning and characterization of mouse extracellular-signal-regulated protein kinase 3 as a unique gene product of 100 kDa. Biochem J 346 Pt 1:169–175.

4. Rousseau J, Klinger S, Rachalski A, Turgeon B, Deleris P, Vigneault E, Poirier-Heon JF, Davoli MA, Mechawar N, El Mestikawy S, Cermakian N, Meloche S. 2010. Targeted inactivation of Mapk4 in mice reveals specific nonredundant functions of Erk3/Erk4 subfamily mitogen-activated protein kinases. Mol Cell Biol 30:5752–5763.

5. Turgeon B, Saba-El-Leil MK, Meloche S. 2000. Cloning and characterization of mouse extracellular-signal-regulated protein kinase 3 as a unique gene product of 100 kDa. Biochem J 346:169–175.

6. Coulombe P, Rodier G, Pelletier S, Pellerin J, Meloche S. 2003. Rapid turnover of extracellular signal-regulated kinase 3 by the ubiquitin-proteasome pathway defines a novel paradigm of mitogen-activated protein kinase regulation during cellular differentiation. Mol Cell Biol 23:4542–4558.

7. Deleris P, Rousseau J, Coulombe P, Rodier G, Tanguay PL, Meloche S. 2008. Activation loop phosphorylation of the atypical MAP kinases ERK3 and ERK4 is required for binding, activation and cytoplasmic relocalization of MK5. J Cell Physiol 217:778–788.

8. De la Mota-Peynado A, Chernoff J, Beeser A. 2011. Identification of the atypical MAPK Erk3 as a novel substrate for p21-activated kinase (Pak) activity. J Biol Chem 286:13603–13611.

9. Deleris P, Trost M, Topisirovic I, Tanguay PL, Borden KL, Thibault P, Meloche S. 2011. Activation loop phosphorylation of ERK3/ERK4 by group I p21-activated kinases (PAKs) defines a novel PAK-ERK3/4-MAPK-activated protein kinase 5 signaling pathway. J Biol Chem 286:6470–6478.

10. Perander M, Al-Mahdi R, Jensen TC, Nunn JA, Kildalsen H, Johansen B, Gabrielsen M, Keyse SM, Seternes OM. 2017. Regulation of atypical MAP kinases ERK3 and ERK4 by the phosphatase DUSP2. Sci Rep 7:43471.

11. Schumacher S, Laass K, Kant S, Shi Y, Visel A, Gruber AD, Kotlyarov A, Gaestel M. 2004. Scaffolding by ERK3 regulates MK5 in development. EMBO J 23:4770–4779.

12. Al-Mahdi R, Babteen N, Thillai K, Holt M, Johansen B, Wetting HL, Seternes OM, Wells CM. 2015. A novel role for atypical MAPK kinase ERK3 in regulating breast cancer cell morphology and migration. Cell Adh Migr 9:483–494.

13. Klinger S, Turgeon B, Levesque K, Wood GA, Aagaard-Tillery KM, Meloche S. 2009. Loss of Erk3 function in mice leads to intrauterine growth restriction, pulmonary immaturity, and neonatal lethality. Proc Natl Acad Sci U S A 106:16710–16715.

14. Kornev AP, Taylor SS. 2010. Defining the conserved internal architecture of a protein kinase. Biochim Biophys Acta 1804:440–444.

15. West DB, Engelhard EK, Adkisson M, Nava AJ, Kirov JV, Cipollone A, Willis B, Rapp J, de Jong PJ, Lloyd KC. 2016. Transcriptome Analysis of Targeted Mouse Mutations Reveals the Topography of Local Changes in Gene Expression. PLoS Genet 12:e1005691.

16. Bradley A, Anastassiadis K, Ayadi A, Battey JF, Bell C, Birling MC, Bottomley J, Brown SD, Burger A, Bult CJ, Bushell W, Collins FS, Desaintes C, Doe B, Economides A, Eppig JT, Finnell RH, Fletcher C, Fray M, Frendewey D, Friedel RH, Grosveld FG, Hansen J, Herault Y, Hicks G, Horlein A, Houghton R, Hrabe de Angelis M, Huylebroeck D, Iyer V, de Jong PJ, Kadin JA, Kaloff C, Kennedy K, Koutsourakis M, Lloyd KC, Marschall S, Mason J, McKerlie C, McLeod MP, von Melchner H, Moore M, Mujica AO, Nagy A, Nefedov M, Nutter LM, Pavlovic G, Peterson JL, Pollock J, Ramirez-Solis R, et al. 2012. The mammalian gene function resource: the International Knockout Mouse Consortium. Mamm Genome 23:580–586.

17. Jokela H, Hakkarainen J, Katkanaho L, Pakarinen P, Ruohonen ST, Tena-Sempere M, Zhang FP, Poutanen M. 2017. Deleting the mouse Hsd17b1 gene results in a hypomorphic Naglu allele and a phenotype mimicking a lysosomal storage disease. Sci Rep 7:16406.

18. Maguire S, Estabel J, Ingham N, Pearson S, Ryder E, Carragher DM, Walker N, Sanger MGPSaPT, Bussell J, Chan WI, Keane TM, Adams DJ, Scudamore CL, Lelliott CJ, Ramirez-Solis R, Karp NA, Steel KP, White JK, Gerdin AK. 2014. Targeting of Slc25a21 is associated with orofacial defects and otitis media due to disrupted expression of a neighbouring gene. PLoS One 9:e91807.

19. Olson EN, Arnold HH, Rigby PW, Wold BJ. 1996. Know your neighbors: three phenotypes in null mutants of the myogenic bHLH gene MRF4. Cell 85:1–4.

20. Pan Y, Zhang L, Liu Q, Li Y, Guo H, Peng Y, Peng H, Tang B, Hu Z, Zhao J, Xia K, Li JD. 2016. Insertion of a knockout-first cassette in Ampd1 gene leads to neonatal death by disruption of neighboring genes expression. Sci Rep 6:35970.

21. Scacheri PC, Crabtree JS, Novotny EA, Garrett-Beal L, Chen A, Edgemon KA, Marx SJ, Spiegel AM, Chandrasekharappa SC, Collins FS. 2001. Bidirectional transcriptional activity of PGK-neomycin and unexpected embryonic lethality in heterozygote chimeric knockout mice. Genesis 30:259–263.

22. Ashburner M, Ball CA, Blake JA, Botstein D, Butler H, Cherry JM, Davis AP, Dolinski K, Dwight SS, Eppig JT, Harris MA, Hill DP, Issel-Tarver L, Kasarskis A, Lewis S, Matese JC, Richardson JE, Ringwald M, Rubin GM, Sherlock G. 2000. Gene ontology: tool for the unification of biology. The Gene Ontology Consortium. Nat Genet 25:25–29.

23. Huang da W, Sherman BT, Lempicki RA. 2009. Systematic and integrative analysis of large gene lists using DAVID bioinformatics resources. Nat Protoc 4:44–57.

